# Identification of a large cohort of *Enterobacter* jumbo phages with broad host ranges across pathogenic Gammaproteobacteria

**DOI:** 10.1101/2024.10.29.620934

**Authors:** Natasha Kanhirun, Alisha Blanc, Ann Aindow, Millie You, Jyot D. Antani, Pooja Ghatbale, Jesse Leonard, Andrew G. Garcia, Kristin Nghiem, Katrine Whiteson, Robert T. Schooley, Paul E. Turner, Joe Pogliano, Ana G. Cobián Güemes, David T. Pride

## Abstract

ESKAPE pathogens cause most hospital-acquired infections globally and often carry antibiotic resistance. Many of them have been the target of bacteriophage therapies. *Enterobacter* is an ESKAPE pathogen but is less frequently a target for phage therapy due to a relative lack of available phages. We isolated eight jumbo phages with genomes ranging from 223 to 366 kbp targeting *Enterobacter* spp. and found that they belonged to separate phage clades. Six of them formed nucleus-like structures confirmed by DAPI-staining, and were phylogenetically related to *Chimalliviridae*. Two jumbo phages did not form nucleus-like structures and did not cluster with *Chimalliviridae*. Although these jumbo phages were found on *Enterobacter*, many were closely related to phages with non-*Enterobacter* hosts. To test whether these phages may have had expanded host ranges, we examined 14 pathogenic Gammaproteobacteria and found that these phages were capable of creating plaques on 8 of them. These species included *Escherichia coli*, *Klebsiella aerogenes*, *Serratia marcescens*, *Salmonella* spp., *Shigella* spp., *Providencia* spp., *Citrobacter* spp., and *Cronobacter sakazakii*. We verified that there was phage amplification in these microbes rather than lysis from without by performing qPCR to confirm DNA replication in each species. Phages typically have narrow host ranges, a benefit for microbiome-sparing compared to antibiotics. However, the broad host ranges of these Gammaproteobacteria jumbo phages suggests that not all phages have the same risk/benefit ratios. While this broad range could aid their development as antibiotic alternatives, further study is needed to assess potential microbiome disruption.

**Significance:** With the growing threat of antibiotic resistant bacteria, alternative treatments like bacteriophages have emerged. Bacteriophages typically have narrow host ranges, a disadvantage compared to antibiotics. We discovered eight jumbo phages that kill antibiotic resistant *Enterobacter*, but were diverse phylogenetically. Six of them formed nucleus-like structures and were members of the *Chimalliviridae* family and the other two did not form nucleus-like structures. We confirmed that each phage had broad host ranges capable of lysing at least eight different human Gammaproteobacteria including pathogens such as *E. coli*, *K. aerogenes*, and *S. marcescens*. By identifying broad host range jumbo phages that attack pathogens, we may have identified phages with spectrums of activity more similar to antibiotics than have been traditionally attributed to phages.

## Introduction

Since the discovery of antibiotics^1^, their use has become widespread in the medical^2^ and agricultural^3^ world. The dependence of antibiotics as the regimen for treating bacterial infections combined with the overuse of antibiotics has resulted in the widespread acquisition of antimicrobial resistance (AMR) and multidrug resistance (MDR)^4^. This ongoing and increasing threat is now a global public health issue best exemplified by the presence of ESKAPE pathogens (*Enterococcus faecium, Staphylococcus aureus, Klebsiella pneumoniae, Acinetobacter baumannii, Pseudomonas aeruginosa,* and *Enterobacter* spp.), which are the leading cause of hospital-acquired infections^5^. These pathogens have become a major challenge in modern-day health care as their resistance often forces alternative treatment protocols^6^.

Among the ESKAPE pathogens, *Enterobacter* spp. have a particular propensity towards antimicrobial resistance. They frequently are known to acquire resistance via plasmids that confer resistance to cephalosporins such as ESBLs (Extended Spectrum Beta-Lactamases) and carbapenems^7^. *Enterobacter* is best known for its expression of AmpC beta-lactamases that confer resistance to a number of cephalosporins and other beta-lactam antibiotics, which render them difficult to treat with antibiotics other than carbapenems^8^. The presence of these antibiotic resistance factors renders *Enterobacter spp.* one of the top targets for the development of alternative therapies to antibiotics.

One promising approach in combating AMR pathogens involves viruses called bacteriophages (phages) that infect and kill bacteria but do not infect humans^9^. Using phages in this manner in a process called phage therapy has been successful in treating AMR bacterial infections^10^. Phages often have high specificity for their host bacteria, which results in relatively limited host ranges for most of them^11^. This is in direct contrast to most antibiotics, which are able to kill bacteria across a spectrum of an entire species or sometimes more generally^12^. The recent rise in global antibiotic resistance combined with the limited host ranges in phages has led to increasing interest in developing phages with broader host ranges against AMR bacteria^13^.

Jumbo phages are tailed bacteriophages with genome sizes ≥ 200 kbp. The large genomes of these phages encode complex machinery for intracellular organization, enzymes that lyse bacterial cell walls^14^, and receptors that often confer wider host ranges^14^. Jumbo phages within the *Chimalliviridae* family^15^, form a proteinaceous “phage nucleus” structure during replication that separates transcription from translation, and protects the phage genome from host defense factors such as CRISPR-Cas systems and Restriction-Modification systems^16–18^.

In this study, we identified 8 unique jumbo phages that infected *Enterobacter* hosts. Our goals were to characterize these jumbo phages to decipher whether they form a phage nucleus, determine how they relate to other known jumbo phages, identify their host ranges across Enterobacter isolates, and to decipher whether they were capable of killing other unrelated pathogenic Gammaproteobacteria.

## Results

### Isolation of 8 jumbo phages on an *Enterobacter cloacae* host

Phages Silp (Si), Carethers (Ca), Rett (Re), Arina (Ar), Sayo (Sa), Chasing Life (CL), Winchester Ellie (WE), and Big Picture (BP) were isolated from filtered wastewater acquired on the University of California San Diego (UCSD) campus^19^. Each of these phages were isolated using *Enterobacter cloacae* bacterial hosts EBC3 (SAMN44407723) (Si, Ar, Re) and EBC7 (SAMN44407724) (Ca, Sa, CL, WE, BP) using a new jumbo phage isolation protocol. This protocol differed from the typical phage isolation protocols in the following ways: the isolation source (wastewater) was not filtrated to avoid separating potential jumbo phages and all resulting plaques were purified.

### Genomic characterization of phages

We sequenced each of the genomes of these jumbo phages using previously described protocols^20^. Each of the phages had double-stranded DNA genomes that ranged from 223,602 to 366,169 bp in size, meeting the minimum criteria for classification as jumbo phages^14^ (**Table 1**). We developed a phylogeny including each of these 8 phages as well as 80 previously sequenced dsDNA jumbo phage genomes to identify whether these new jumbo phages were related to any of those previously sequenced genomes (**Fig. 1, Table S1**). We identified 4 separate clades in the Pseudomonadota host group that included our viruses, with the first clade including Si, Ca, Re, and Re. Each of these four phages had genome sizes around 244 kbp. The second clade included Sa, with a genome size of 223 kbp, and CL, with a genome size of 276 kbp. The third clade included WE with a genome size of 259 kbp. BP, clustered the furthest away from the majority of the phages in a separate fourth clade, and had the largest genome size of 366 kbp.

**Figure 1.**
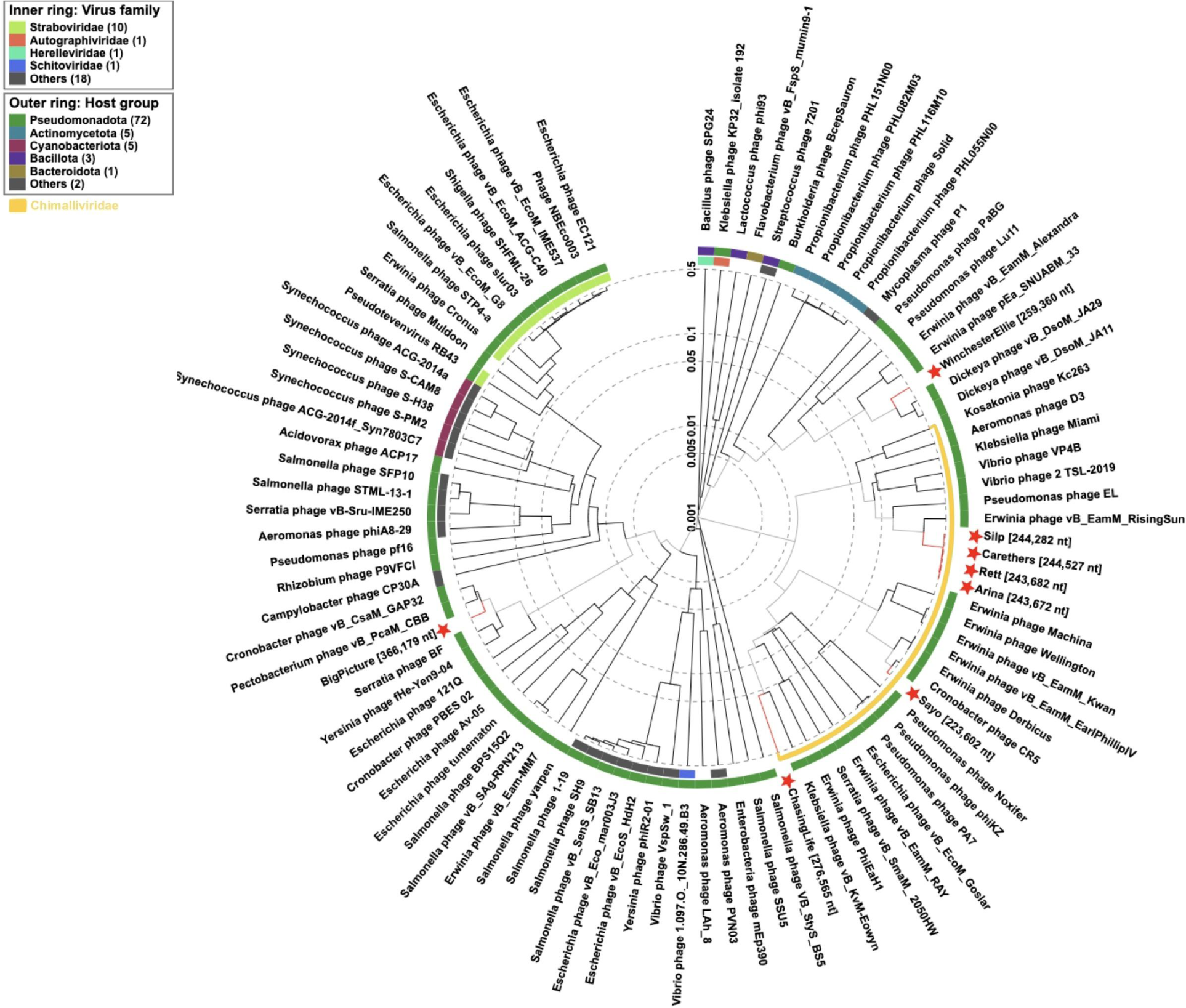
Phage proteomic tree showing the relationship between jumbo phages. The dendrogram represents proteome-wide similarity relationships among the 8 phages and 88 prokaryotic dsDNA virus genomes. Branches are colored black (dsDNA virus genome) or red (the jumbo phage genome). Branch lengths are indicated using a logarithmic scale. For the outside colored rings, the inner ring represents the viral family classifications and the outer colored ring represents the taxonomic groups of known hosts. The *Chimalliviridae* clade is highlighted in yellow. The 8 new *Enterobacter* jumbo phages are highlighted by red stars.

Genome sequence alignments to the phages’ closest related genomes showed that Si, Ca, Re, and Re are closely related to *Erwinia* phage RisingSun^21^ (**Fig. S1A**). The closest matches for Sa, CL, WE, and BP were *Cronobacter* phage CR5^22^ (**Fig. S1B**), *Klebsiella* phage Eowyn^23^ (**Fig. S1C**), *Dickeya* phage JA29^24^ (**Fig. S1D**), and *Pectobacterium* phage CBB^25^ (**Fig. S1E**), respectively. Although these phages were isolated on *Enterobacter* hosts, they were more closely related to other non-*Enterobacter* Gammaproteobacteria phages.

### Morphological characterization and relationships to related phages

We performed transmission electron microscopy (TEM) for all 8 phages to identify their morphological characteristics (**Fig. 2A-H**). Each of the 8 phages shared common characteristics of a jumbo bacteriophages and exceeded 200 nm in length. Their displays of an icosahedral head, contractile tail, and thin tail fibers attached to a baseplate verified their myovirus-like morphologies^26^. Some phages showed crossed striations on the tail sheaths leading up to their prominent base plates and others depicted hexagonal outlines on their icosahedral heads (**Fig. 2A-H**).

**Figure 2.**
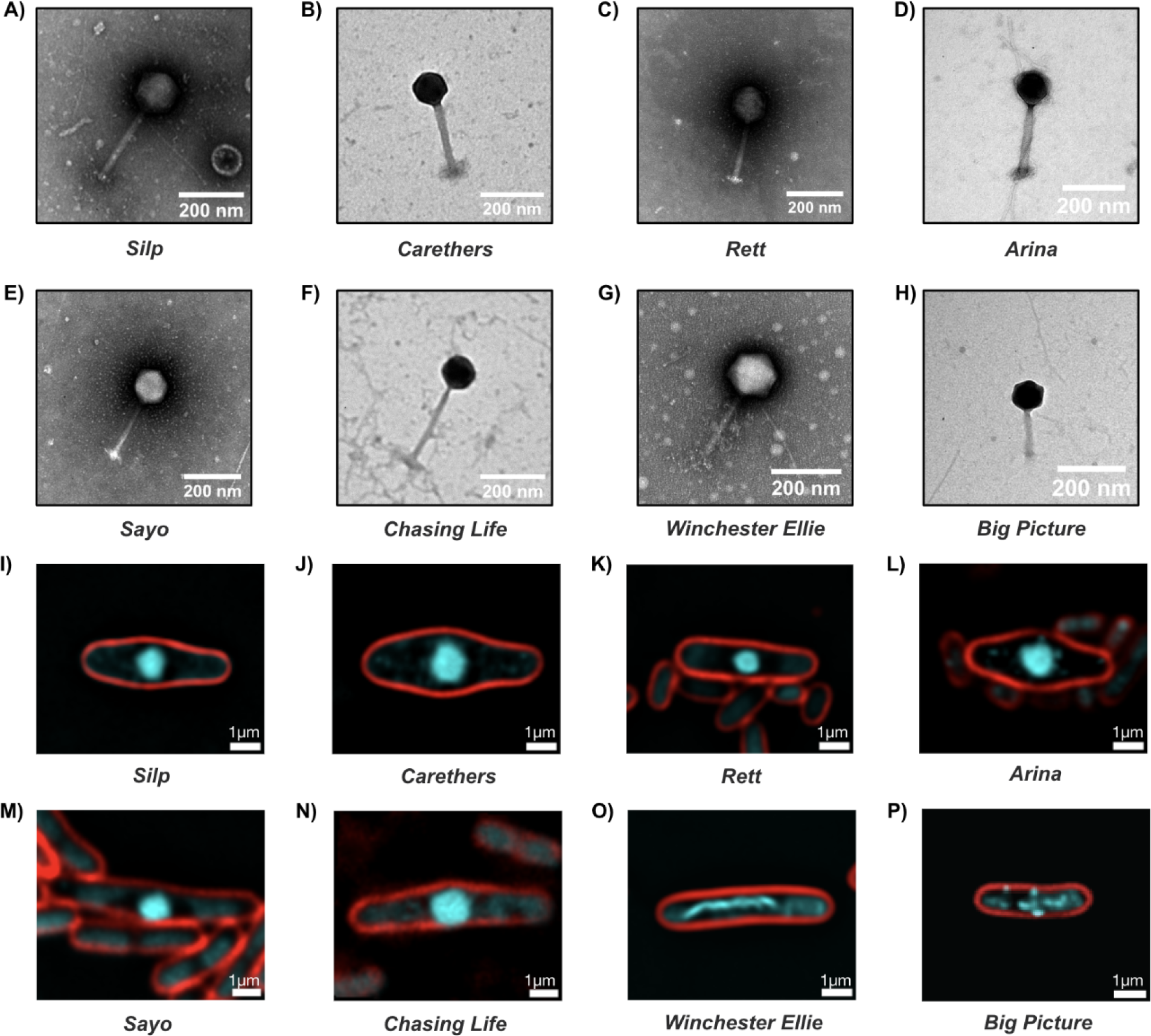
Jumbo phage morphology and infection phenotype. A-H) Transmission Electron Micrograph (TEM) of each phage. Phage dimensions are measured using the image capture software ImageJ. Scale bars are 200 nm. I-P) Fluorescence microscopy of *Enterobacter cloacae* cells infected with jumbo phages. Cells were imaged at 90 minutes post-infection(mpi). The cell membrane is stained with FM4-64 (red), and DNA is stained with DAPI (cyan). The phage nucleus is seen centered in the cell (I-N). Scale bars are 1μm. A,I) Silp. B,J) Carethers. C,K) Rett. D,L) Arina. E,M) Sayo. F,N) Chasing Life. G,O) Winchester Ellie. H,P) Big Picture.

### Nucleus formation of jumbo phages

Out of the 8 phages, 6 phages (all except WE and BP) reside within the chimallin (ChmA)-encoding monophyletic clade called *Chimalliviridae*^15^ (**Fig. 1**). We performed live-cell fluorescence microscopy^15^ on each of these phages infecting their host bacteria to determine whether they formed a phage nucleus. Phage nucleus formation was observed in EBC7 cells infected with each phage and were captured at 90 minutes post-infection (mpi) with DAPI staining (**Fig. 2**). These images confirmed that Si, Ca, Re, Ar, Sa, and CL formed a phage nucleus within the bacterial cell during phage replication (**Fig. 2I-N**), while WE and BP did not (**Fig. 2O,P**), as was predicted by their position on the phylogenies (**Fig. 1**).

### Jumbo phages adsorption time and lysis time

To discern how these jumbo phages were interacting with host *Enterobacter* cells, we performed microscopic adsorption assays by fluorescently labeling each of the phages to calculate mean dwell times – the average time that a phage interacts with bacterial cells. We recently demonstrated that mean dwell times correlate with phage adsorption rates measured via traditional bulk methods.^27^ The distributions of mean dwell times for each of the phages was between 1 and 1.5 seconds (**Fig. S3A)**. Next, we determined the lysis time of single EBC7 cells infected with each jumbo phage using time-lapse phase-contrast microscopy. We calculated the average single-cell lysis time for each experiment (**Fig. S3B**). We observed lysis times ranging from 100 to 250 minutes, which were consistent with previously reported lysis times for other jumbo phages.^28–32^

### Jumbo phages are active against a number of different *Enterobacter* spp. Isolates

To discern whether the 8 phages we isolated have relatively broad host ranges across *Enterobacter cloacae* and other *Enterobacter* spp., we tested each of them against a collection of clinical isolates (**Fig. 3A**). Hosts that lysed upon mitomycin C treatment, indicative of the presence of prophages in their genome, were excluded from this screen, resulting in 76 *Enterobacter* strains (data not shown). At least one phage inhibited the growth of 62 out of the 76 (82%) isolates (**Fig. 3A**), indicating that there was relatively broad host range for these phages across this *Enterobacter* isolate collection. Considering that *Enterobacter cloacae* is the most clinically relevant of the *Enterobacters*, we found that 100% of *E. cloacae* isolates could be lysed by our jumbo phage collection (particularly phages Sa and Si), a number that rivals the efficacy of typical antibiotics^33^. These data indicate that these phages had relatively broad host ranges across *Enterobacter* isolates and suggest that their antimicrobial activity might approach that of conventional antibiotics.

**Figure 3.**
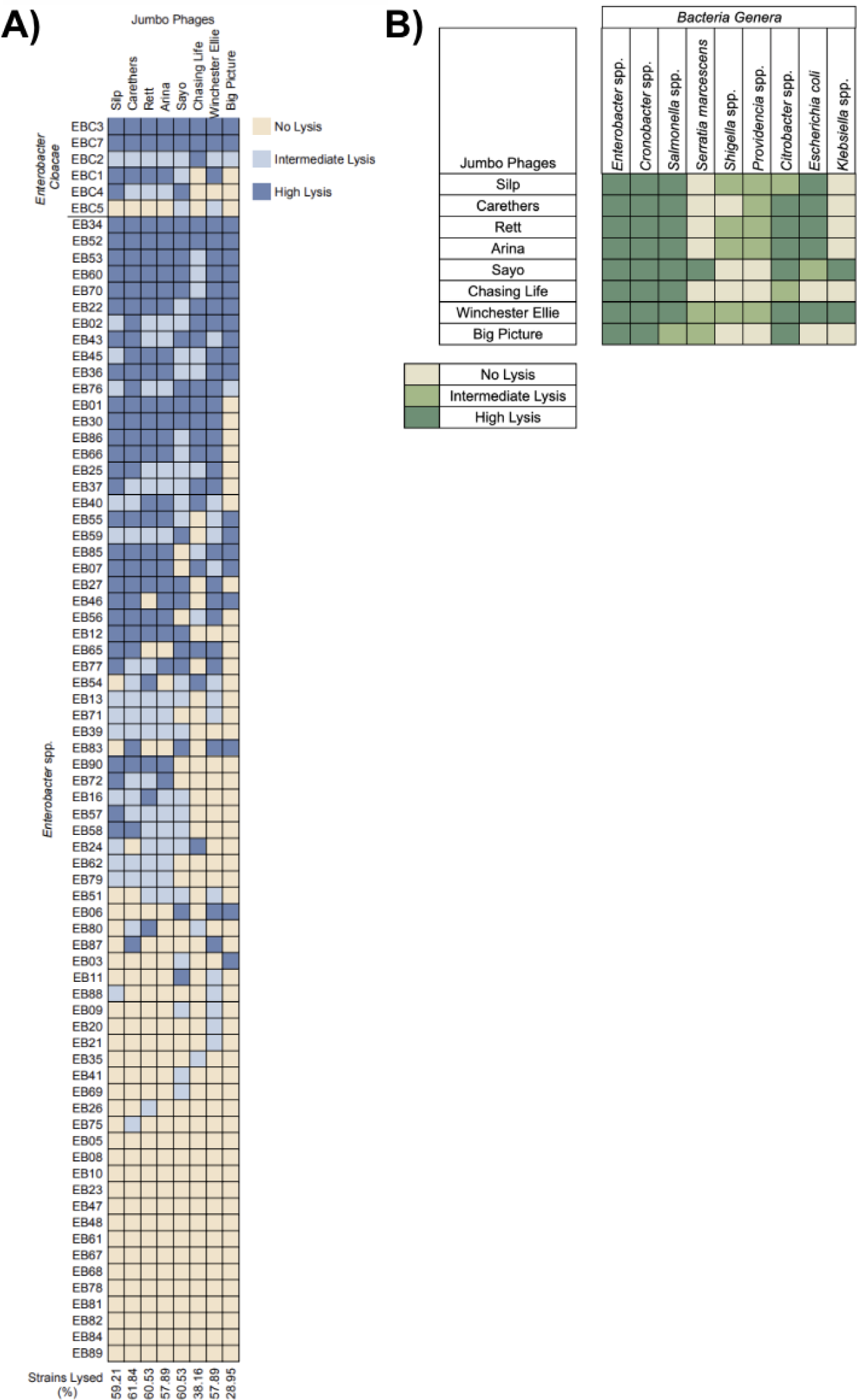
Jumbo phage host range. A) *Enterobacter* host range. Jumbo phage activity was evaluated via spot titer in solid media. Each phage was tested against 70 *Enterobacter* spp. clinical strains and 6 *Enterobacter cloacae* clinical strains. Percentage of strains lysed for each phage is shown. B) Jumbo phages extended host range. Summarized host range of all tested combinations of jumbo phages and bacteria that had lysis. Phages were tested against 14 genera of bacteria, 8 of which were lysed by at least one phage. The highest lysis for any strain is displayed for each phage.

### Jumbo phages are active against other Gammaproteobacteria

Analyses of the sequences and phylogenetic positioning of these jumbo phages suggested that many may infect hosts other than *Enterobacter* spp. (**Fig. S1**). We hypothesized that these phages either have definitive hosts other than *Enterobacter spp.*, or they may be broad host range phages, which have not been frequently described in the literature. Having access to rather large collections of clinical isolates of human pathogens, we investigated whether these 8 jumbo phages isolated on *Enterobacter* hosts were capable of killing other similar bacteria. We focused primarily on Gammaproteobacteria and Betaproteobacteria, as those are related clades and have high propensities to cause illnesses in humans^34^. We first examined whether the phages were capable of lysing *Cronobacter*, which is a pathogen closely related to *Enterobacter*. We found that all 8 phages were capable of killing all 16 tested isolates of *Cronobacter sakazakii* (**Fig. 3B, Fig.S3A**). We also found that the phages collectively were capable of killing 6/7 isolates of *Citrobacter* spp. **(Fig. S3B),** 6/20 isolates of *Escherichia coli* **(Fig. S3C),** 3/4 isolates of *Klebsiella aerogenes* **(Fig. S3D),** 2/19 isolates of *Providencia spp* **(Fig. S3E),** 11/21 isolates of *Salmonella* spp. **(Fig. S3F)**, 14/15 isolates of *Serratia marcescens* **(Fig. S3G)**, and 4/20 isolates of *Shigella* spp. **(Fig. S3H)**. While these data strongly suggested broad host ranges amongst these 8 jumbo phages, no lytic potential was observed for some bacterial species. For example, we tested these phages against *Morganella* spp., *Acinetobacter* spp., *Stenotrophomonas maltophila*, *Burkholderia* spp., *Achromobacter* spp., as well as *Pseudomonas* spp., and found no evidence of lysis in any of these species (**Figure S4**). These data suggested that while these jumbo phages appear to have broad host ranges, the observed lysis could be due to lysis from without^35^ rather than phage replication.

### Jumbo phages actively replicate in other host bacteria

To discern whether there was active replication of the 8 jumbo phages within each of the different bacterial species with evidence of lysis, we performed qPCR analysis to determine whether phage DNA was replicating in these hosts (**Fig. 4**). This would help us decipher whether bacterial lysis was occurring without active DNA replication within the hosts. We performed these experiments in each of the bacterial hosts with each of the 8 separate phages at both time 0 and at 18 hours of phage infection. We then extracted DNA at each time-point (0 and 18 hours) and performed qPCR with phage specific primers to quantify the relative abundances of each phage (**Fig. 4**). We found that, in many instances, a phage actively replicated in a host when it was observed to create plaques in its host range assay. Conversely, turbid plaques or the absence of lysis often corresponded to a lack of active replication in the tested bacteria.

**Figure 4.**
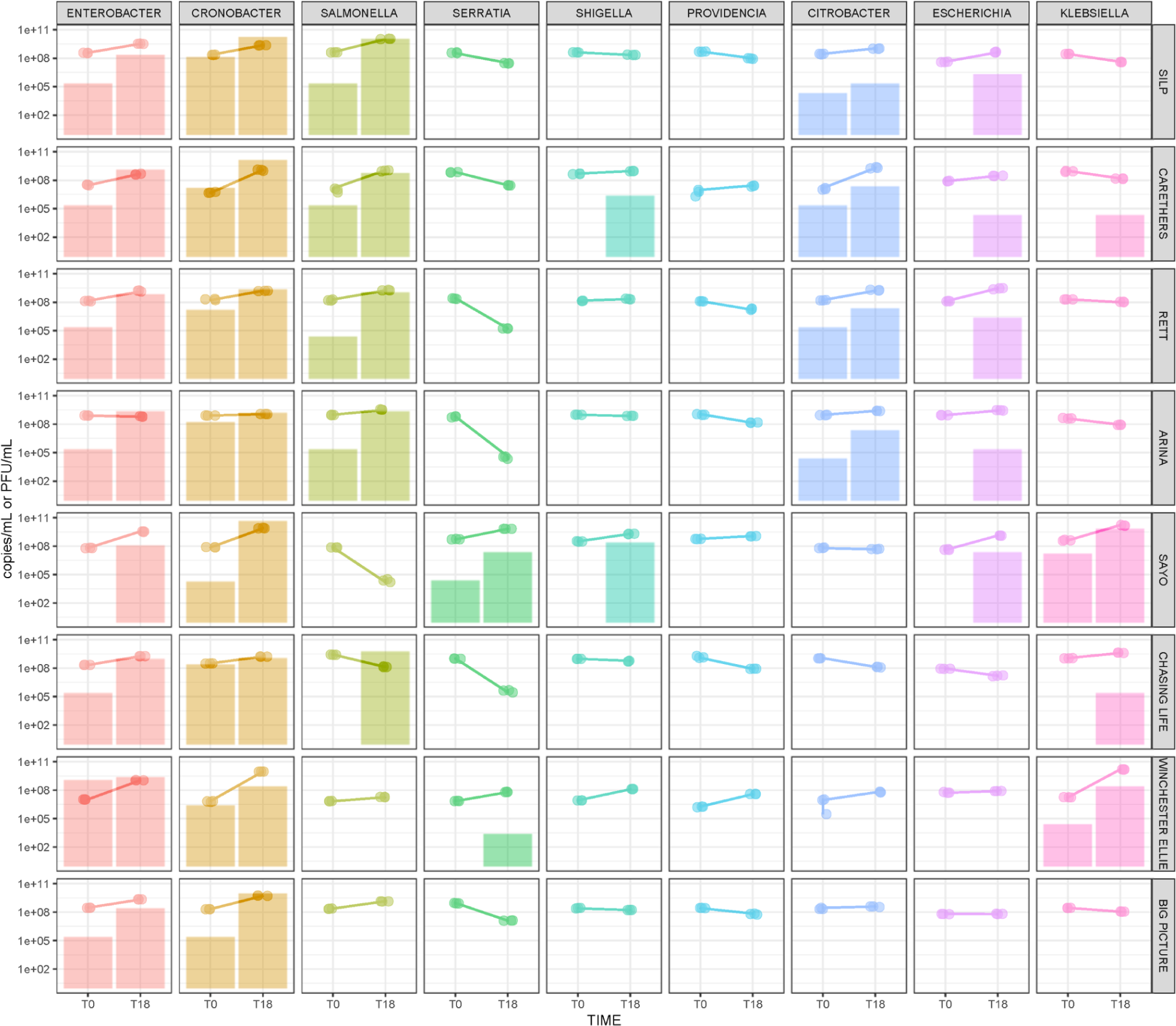
Replication in other hosts by qPCR and phage titer. Phage DNA copies/mL at T0 and T18 are shown with connecting dot plots. Phage titer at T0 and T18 is shown by bar plots. Measurements were obtained for each phage against all 9 bacteria genera representative strains.

## Discussion

When targeting human pathogens, the narrow host range of a bacteriophage is considered both a significant strength and a weakness. It is viewed as a weakness because it limits the ability to use phages as therapeutic options against pathogens, and was likely the reason that phages were replaced by antibiotics as the therapeutics of choice in Western medicine^36^. More recently, narrow host range has been viewed as a strength because of its microbiome-sparing effects compared to conventional antibiotics^37^, which can limit long-term deleterious effects observed with antibiotics. Our work characterizing jumbo phages with broad host ranges across Gammaproteobacteria does not fit into the existing paradigm for phage behavior^29,36^, particularly as pertaining to the use of these phages as therapeutic agents. Thus, the discovery of these novel jumbo phages necessitates that we alter our canonical understanding of phage behavior and investigate the consequences of phage treatment on both pathogens and the rest of the microbiome.

The host ranges of jumbo bacteriophages are often broader due to their larger genome sizes, possibly consisting of additional host lysis and DNA replication genes^38^. We found that our jumbo phages have host ranges across 8 additional Gammaproteobacteria, one of which is not included in the *Enterobacteriaceae* family. We were limited in our tests by the strains that were available to us for testing. It is possible that these phages have broader host ranges across additional Gammaproteobacteria which were not tested because they were not available to us. Particularly because we found that some isolates within a species were susceptible and others were not, susceptibility to these phages is complicated, and testing additional isolates may reveal an even broader host range. To rule out bacterial lysis without active replication of the phages, we confirmed activity of the phages using both plaque assays and qPCR.

Six out of the eight jumbo phages isolated were members of the *Chimalliviridae* family, as evidenced by their phylogenetic position (**Fig. 1**) and ability to form a phage nucleus (**Fig. 2I-N**)^16^. These nucleus-forming phages exhibit different levels of host genome degradation and nucleus centering, two hallmarks of *Chimalliviridae*. However, two of our jumbo phages were from different clades and do not form a phage nucleus. Fluorescence microscopy of these infections with DAPI staining showed one or more regions of high DNA density. In the case of phage WE, DNA appeared to form a filamentous structure (**Fig. 2O**). It is possible that these two phages replicate by a different, unknown mechanism that involves intracellular organization of phage genome or virion components. As the ability to form a nucleus is an advantage against host defense systems, this complex replication pathway should be studied further.

The broad host ranges of these jumbo phages may be advantageous for phage therapy^39^. A broader host range may reduce the risk of treatment failures^40,41^. It also encourages the production of standardized phage cocktails, those of which would only need a small number of phages to cover a wide range of bacterial strains^42^. Lastly, broad host range phages may be preferable in treating mixed bacterial infections as one phage could be sufficient in killing multiple species instead of finding a therapeutic phage for each^42^.

This work highlights the characterization of 8 novel jumbo phages that were isolated on an *Enterobacter* host but have the ability to infect at least 8 other Gammaproteobacteria. Six of these phages undergo replication by forming a nucleus to protect its DNA against host defense factors, and the remaining two do not form a nucleus but may be adopting other mechanisms during their lytic replication cycle. Many of these attributes promote their use in phage therapy to kill multiple genera of bacteria in AMR infections. Further studies should be directed towards understanding their replication and host range mechanisms, and their impacts on the microbiome to build additional confidence in their clinical use.

## Methods

### Jumbo phage isolation and purification

All phages were isolated from the UCSD campus wastewater obtained from the UCSD Knight Lab^19^. Wastewater was centrifuged at 3,000 x g for 10 minutes and the supernatant was collected, then centrifuged at 4,696 × g for 55 minutes^20^. The supernatant was discarded and the pellet was resuspended in 10 mL of SM buffer and gently vortexed. An overnight culture of *Enterobacter cloacae* EBC7 was diluted to 0.2 OD_600_, 200 μL of culture was combined with 4 mL of warmed 0.3% LB top agar and poured over 1% LB plates. The plates were dried for 15 minutes near an open flame. A 10 ul spot of filtered wastewater was done on the plates and dried under an open flame for 30 minutes. Plates were incubated upside down overnight at 37°C. Phages formed ∼1 mm plaques on LB top agar (0.3%) over the bacterial lawns. Phage plaques were picked using a pipette tip and gently stabbed repeatedly into an LB plate with 1% agar. This area was then streaked with a loop, changing loops with each streak. Molten 0.3% LB top agar and 100 ul of overnight 0.2 OD_600_ EBC7 culture were combined and poured onto the plate starting from the lowest dilution point to the stabbed area. The top agar was then allowed to sit at room temperature for 15 minutes before being incubated overnight at 37°C. Plaques were purified 3 times.

### Creation of Jumbo Phage Stock

An overnight culture of strain EBC7 was diluted to 0.2 OD_600_. 100uL of the diluted culture was incubated with 50 ul of phage, either a phage plaque mixed with LB or current phage stock, for 15 minutes at room temperature. This mixture was combined with 4 mL of warmed 0.1% top agar (LB broth with 0.1% agar) and poured over 1% solid media agar plate evenly to make a bacterial lawn. The plates were left to settle for 15 minutes and then incubated, without inverting, at 37°C overnight for 18-24 hours. After incubation, the plate was flooded with 5 ml of SM buffer and allowed to soak for about 3 hours. The top agar slush was collected and centrifuged at 3000 rpm for 10 minutes and filtered with a 0.45 μm filter. Stocks were stored at 4°C.

### Jumbo phages sequencing and imaging

Phages were sequenced and annotated **(Table S2 and S3)**, as well as the bacteria host (Supplemental Methods). TEM and live-fluorescence microscopy were performed as previously described (Supplemental Methods)

### Host Range

All bacterial strains were isolated from the UCSD clinical laboratory. For *Enterobacter* isolates **(Table S4)**, an overnight culture was diluted to 0.2 OD_600_. 200uL of culture was combined with 4 mL of warmed 0.3% LB top agar and poured over 1% solid media agar plate and dried for 15 minutes near an open flame **(Figure S5)**. Four ul of undiluted phages were spotted on each lawn and dried under an open flame for 30 minutes. Plates were incubated overnight at 37°C. Plates were imaged and analyzed. For the extended host range solid assays, 14 genera of Gammaproteobacteria isolates were tested **(Figure S4)**. Phage infection was evaluated by spot test of serial dilutions of phages. **(Supplemental Figure S3 and S4)**.

### Phage Replication

Two jumbo phage stocks, T=0 hour incubation and T=18 hour incubation, were created for each of the 8 phages with EBC7 (*Enterobacter*), CR04 (*Cronobacter*), SH13 (*Shigella*), SL22 (*Salmonella*), SR01 (*Serratia*), PR13 (*Providencia*), CB02 (*Citrobacter*), EC507 (*Escherichia coli*), EB14 (*Klebsiella*), and AX05 (*Achromobacter*). Five hundred μL of phage stock was incubated with 100 μL of 0.2 OD_600_ of selected bacterial culture for 15 minutes at 20°C. The mixture was combined with warmed 0.1% LB top agar and poured onto a 1% LB-agar plate which was allowed to sit for 15 minutes at 20°C (T=0). This process was repeated and the second plate (T=18) was incubated for 18 hours at 37°C. The second plate was then flooded with 5 mL of SM buffer and incubated at 20°C for 3 hours. The top agar slush was then transferred and centrifuged for 10 minutes at 4,696 × g. The supernatant was filtered through a 0.22 μm filter. Lysates were titered and 1 mL was allocated for DNA extraction using DNeasy Blood & Tissue Kit (Qiagen catalog number 69504). qPCR was performed in triplicate on 1:10 dilutions of the extracted DNA with TaqMan Real-Time PCR Master Mix, primers targeting ∼100 bp of the capsid protein sequence (**Table S5**). A standard curve was used and genome copies per mLvalues were calculated from CT values **(Figure S6)**. Results were visualized using Rstudio **(Figure 4)**.

### Microscopic phage adsorption assay

Phages were fluorescently labeled with the lysine-binding dye AZDye TFP Ester 568 (VectorLabs FP-1091) as recently described^27^. *E. cloacae* (EBC7) cells from an overnight culture were concentrated 10× via microcentrifugation and 2 μL cells were inoculated on imaging pads (1.5% agarose, 100% LB) in welled microscope slides. Cells were incubated at room temperature for ∼15 min before adding 1 μL of fluorescently labeled phages and sealing the sample with a coverslip. Time-lapse videos of phages were acquired via epifluorescence microscopy using a Nikon Ti-E inverted microscope (100× magnification, 1.4 NA, Texas Red fluorescent filter set). Individual phage trajectories were obtained via particle tracking in MATLAB, and dwell time (time that a phage particle spends interacting with bacteria) was determined for each phage. The mean dwell time for all phages in a single replicate experiment was calculated as area under the survival probability curve for the phage trajectory durations (explained in detail in this reference^27^).

### Microscopic measurements of lysis time

*E. cloacae* cells (5 μL of EBC7 overnight culture) were inoculated on imaging pads (1.5% agarose, 100% LB) in welled microscope slides. Slides were incubated at 30 °C for 1 h in a humid chamber to allow cells to reach exponential growth phase. 10 μL of phage lysate was added to the pads and the infection was allowed to proceed at 30 °C. Time-lapse videos of bacteria were acquired via phase-contrast microscopy using a Nikon Ti-E inverted microscope (100× magnification, 1.4 NA). Time of lysis after infection was calculated for up to the first 20 single cells in each video., in order to calculate mean lysis time for each biological replicate (movie).

---

**Figure S1.**
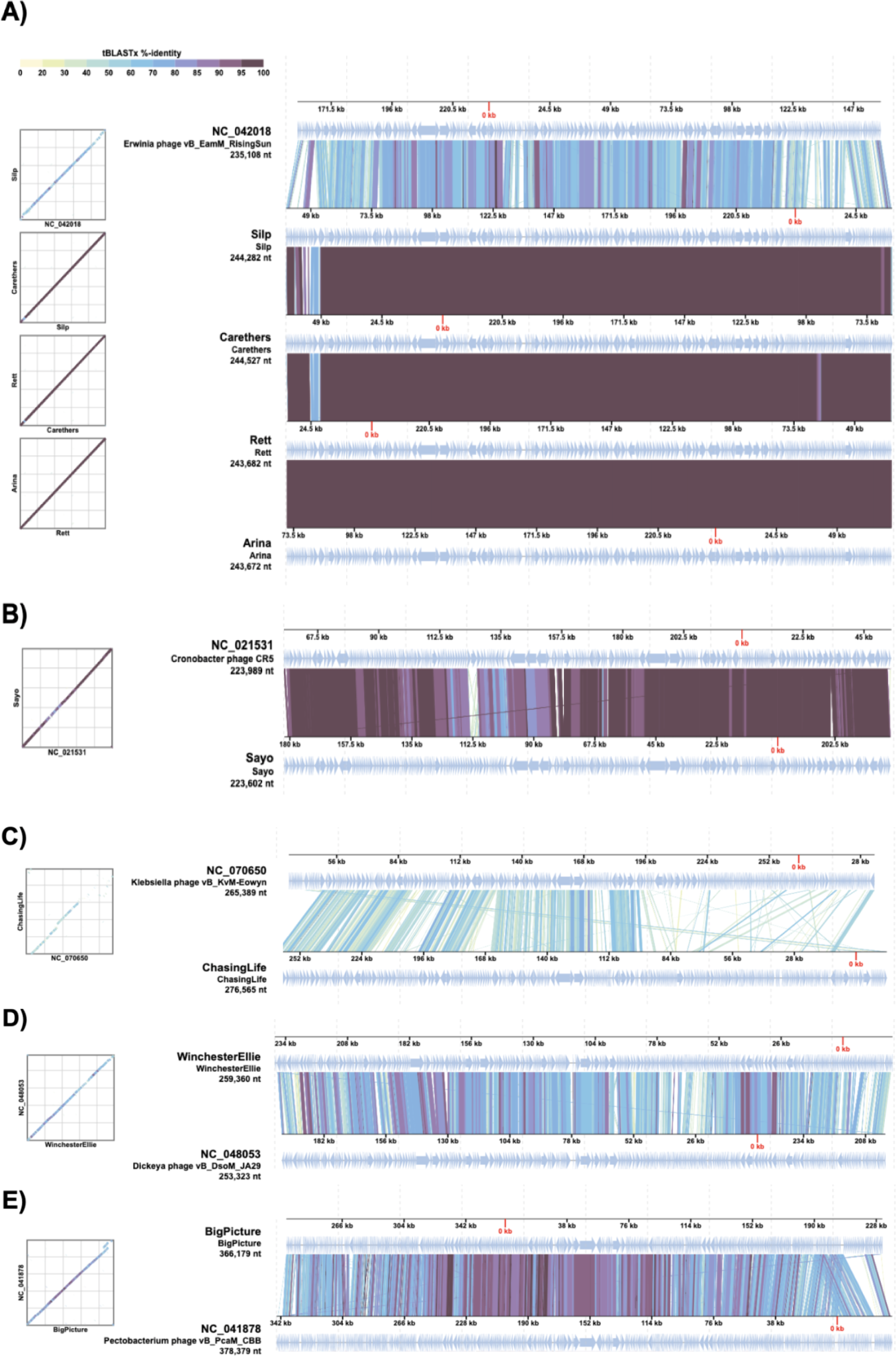
Sequence alignments. The whole sequence of all *Enterobacter* phages against their closest related genome. The colors indicate the tBLASTx Identity percentage between each phage genome and its respective comparison genome. The darkest color indicates portions of the genome that have 100% sequence similarities. **A)** Phages Silp, Carethers, Rett, and Arina compared to *Erwinia* phage Rising Sun. **B)** Phage Sayo compared to *Cronobacter* phage CRS. **C)** Phage Chasing Life compared to *Klebsiella* phage Eowyn. **D)** Phage Winchester Ellie compared to *Dickeya* phage JA29. **E)** Phage Big Picture compared to *Pectobacterium* phage CBB.

**Figure S2.**
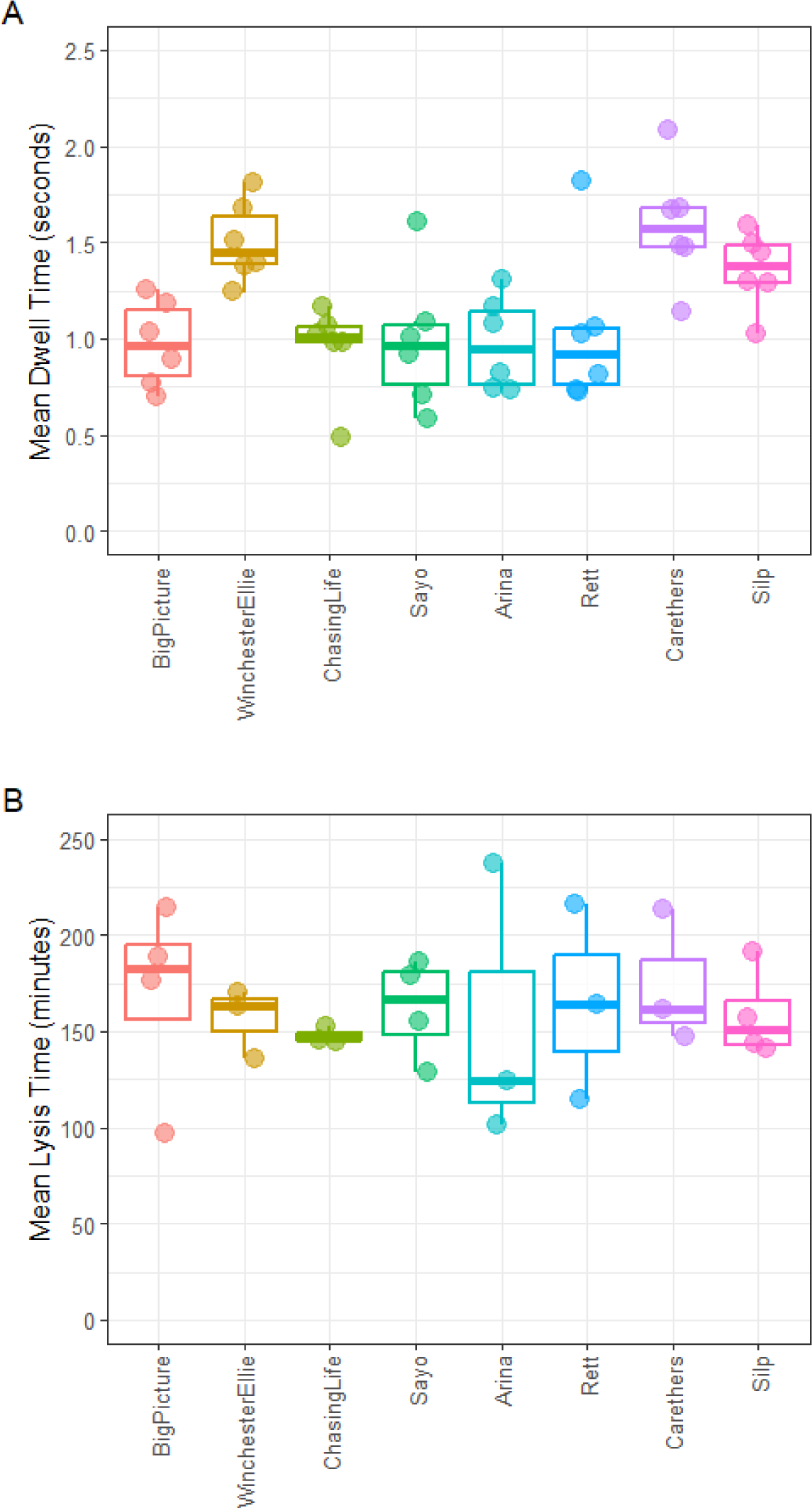
Life cycle characterization of jumbo phages. A) Absorption time B) Replication time. Fluorescently labeled phages were incubated with *Enterobacter cloacae* EBC7 and time lapse videos were obtained in a fluorescence microscope.Single cell trajectories were monitored and absorption time and replication time were obtained from 20 individual cells.

**Figure S3.**
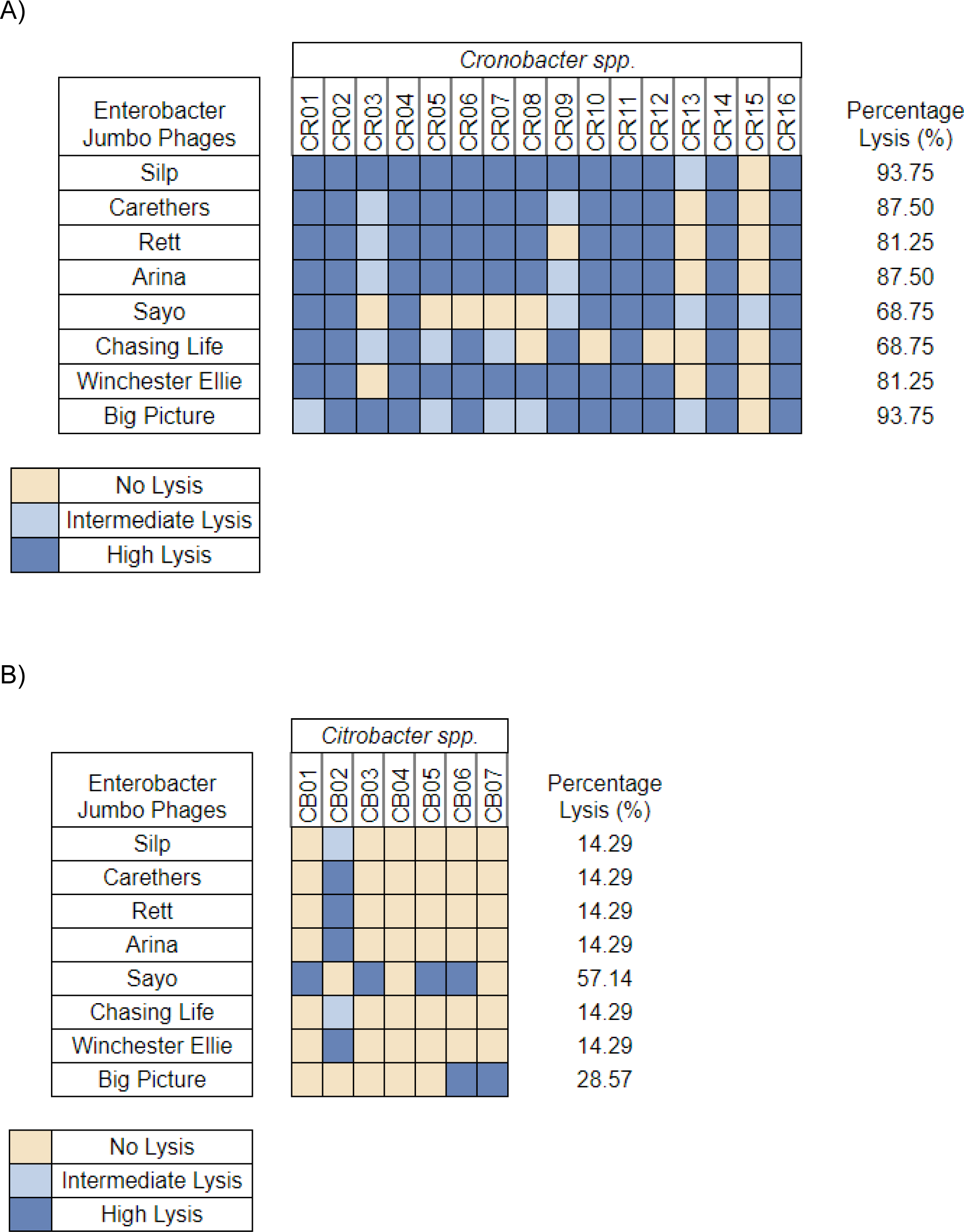
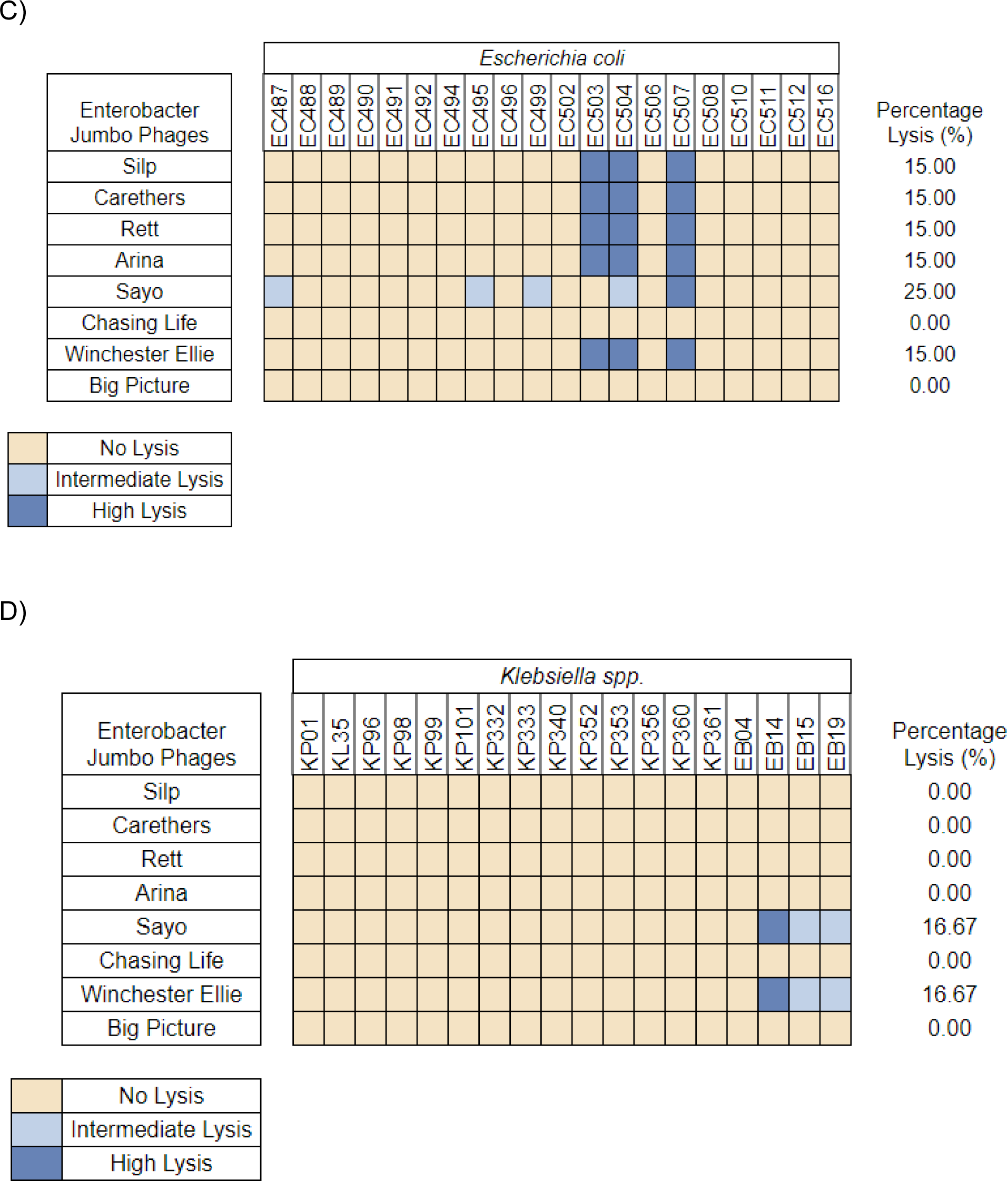
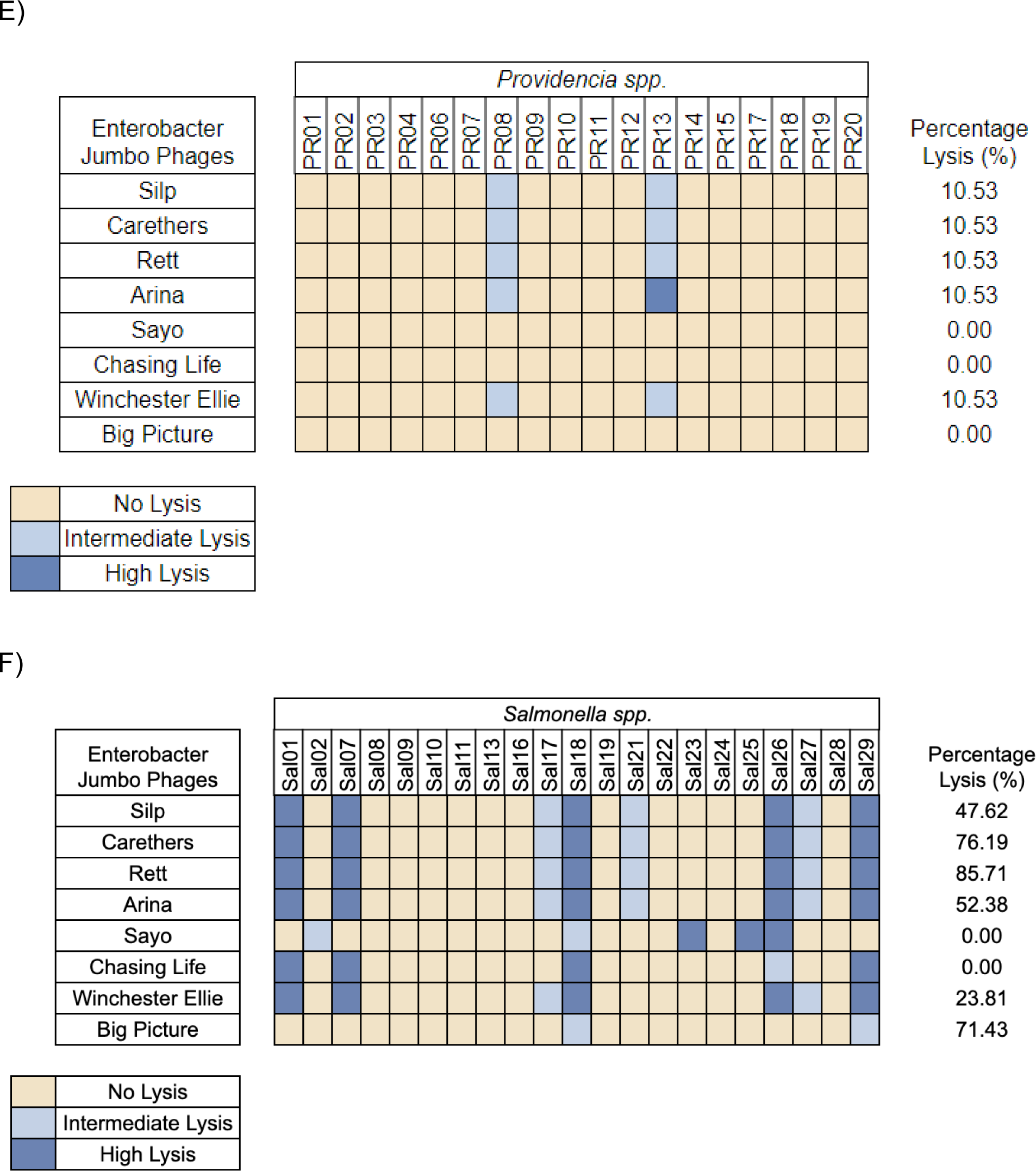
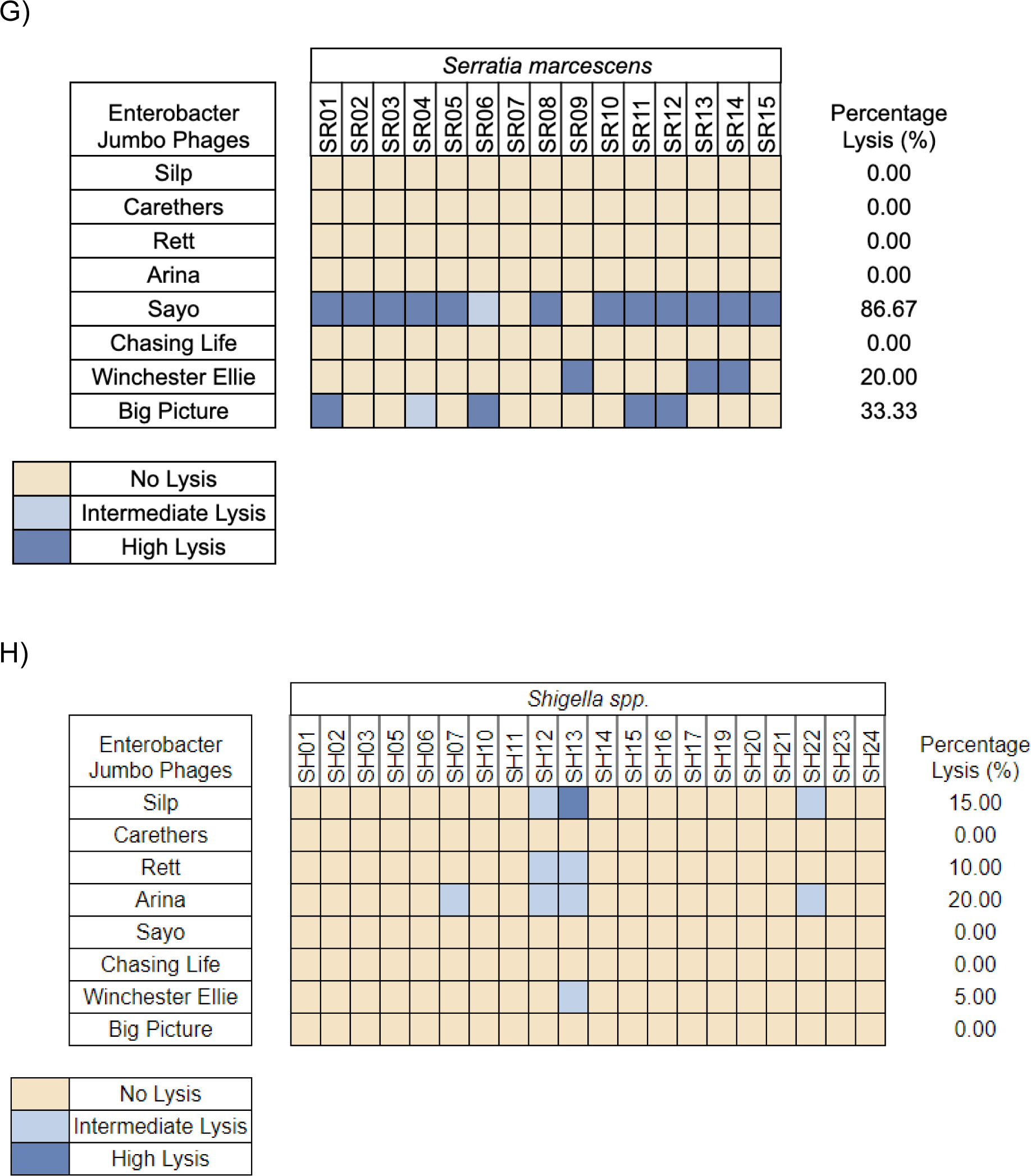
Host range of phages in Gammaproteobacteria. A) Host range of phages in 16 selected isolates of Cronobacter spp. B) 7 selected isolates of *Citrobacter* spp. C) 20 selected isolates of Escherichia coli. D) 18 selected isolates of Klebsiella spp. E) 19 selected isolates of Providencia spp. F) 21 selected isolates of Salmonella spp. G) 15 selected isolates of Serratia marcescens. H) 20 selected isolates of Shigella spp. Phage titers are separated into 3 categorical values. 0 for no lysis, 1-2E3 for intermediate lysis, and values higher than 2E3 for high lysis.

**Figure S4.**
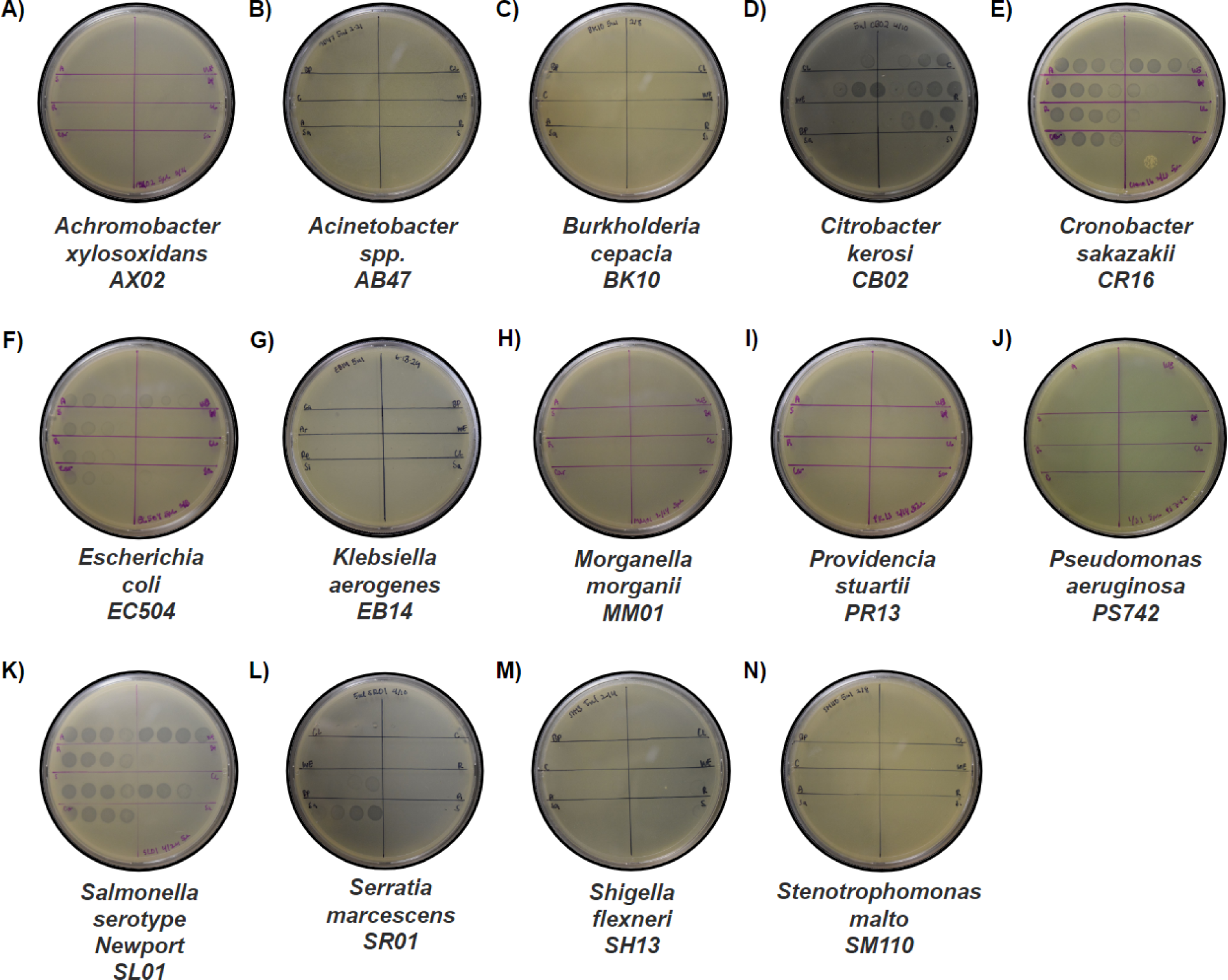
Representative pictures of the extended host range test. Host range assays for all phages against all 14 genera of bacteria. 5 uL spots of all jumbo phages from undiluted to 10E3 dilution and one control spot of LB were spotted. **A)** *Achromobacter xylosoxidans,* strain AX02. **B)** *Acinetobacter* spp., strain AB47 **C)** *Burkholderia cepacia,* strain BK10. **D)** *Citrobacter koseri,* strain CB02. **E)** *Cronobacter sakazakii, strain CR16*. **F)** *Escherichia coli, strain EC504.* **G)** *Klebsiella aerogenes,* strain EB14. **H)** *Morganella morganii,* strain MM01. **I)** *Providencia stuartii,* strain PR13. **J)** *Pseudomonas aeruginosa,* strain PS742. **K)** *Salmonella* spp., strain Group C2C3*, Salmonella serotype Newport.* **L)** *Serratia marcescens,* strain SR01 **M)** *Shigella flexneri,* strain SH13. **N)** *Stenotrophomonas maltophilia,* strain SM110. Phage abbreviations: Silp (S), Carethers (C), Rett (R), Arina (A), Sayo (Sa), Chasing Life (CL), Winchester Ellie (WE), and Big Picture (BP).

**Figure S5.**
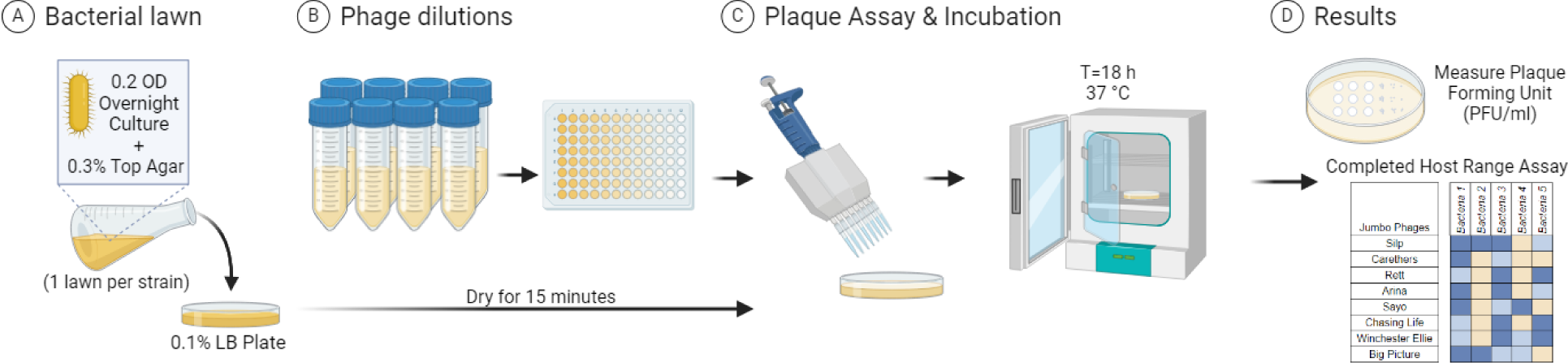
Schematic of phage host range experiment. This protocol was repeated for each phage and each selected bacterial isolate combination.

**Figure S6.**
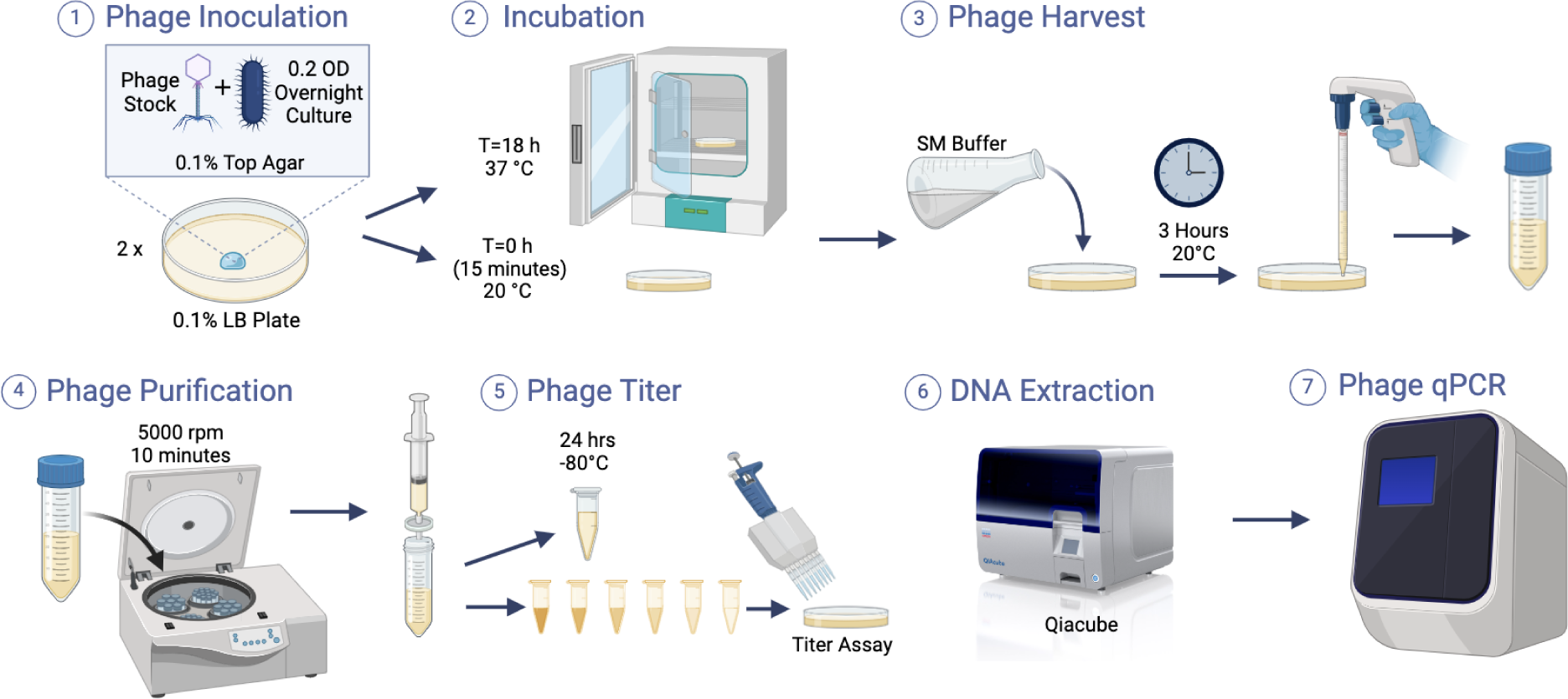
Schematic of phage replication experiment. This protocol was repeated for each phage and each selected bacterial isolate combination.

**Table S1.** Genomes in the phage proteomic tree.

**Table S2.**
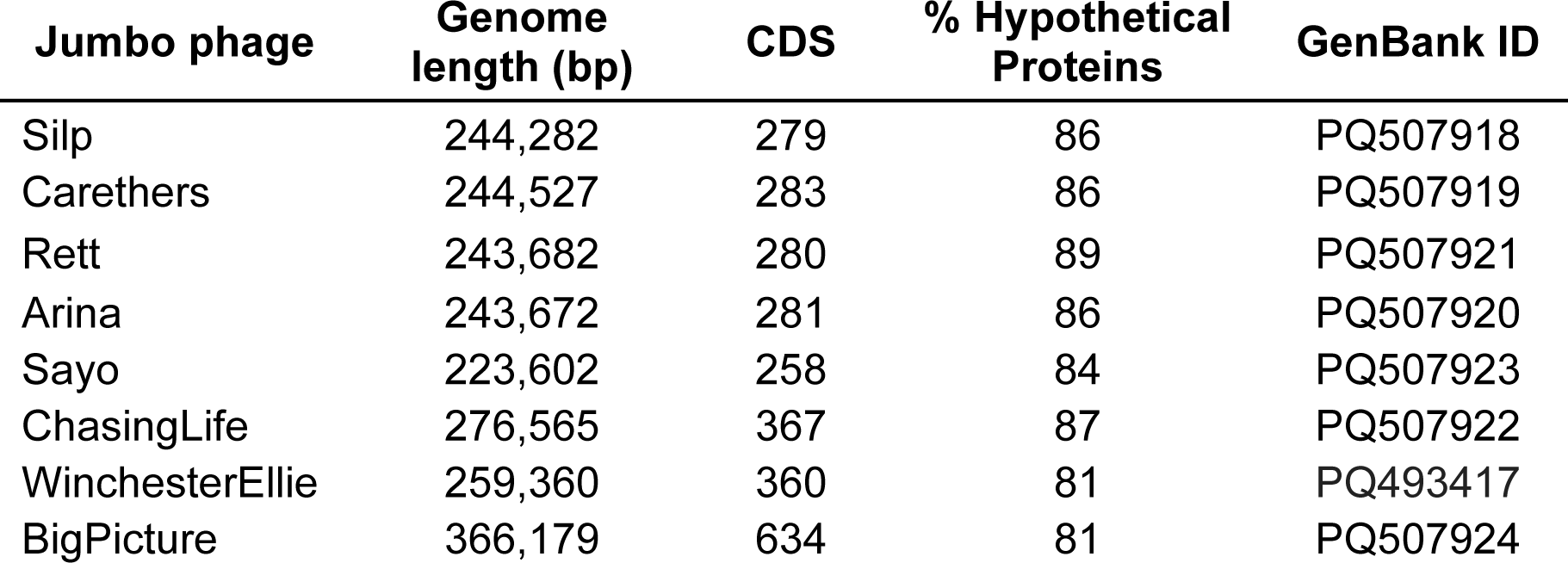
Phage Genome Annotations.

| **Jumbo phage** | **Genome length (bp)** | **CDS** | **% Hypothetical Proteins** | **GenBank ID** |
| --- | --- | --- | --- | --- |
| Silp | 244,282 | 279 | 86 | PQ507918 |
| Carethers | 244,527 | 283 | 86 | PQ507919 |
| Rett | 243,682 | 280 | 89 | PQ507921 |
| Arina | 243,672 | 281 | 86 | PQ507920 |
| Sayo | 223,602 | 258 | 84 | PQ507923 |
| ChasingLife | 276,565 | 367 | 87 | PQ507922 |
| WinchesterEllie | 259,360 | 360 | 81 | PQ493417 |
| BigPicture | 366,179 | 634 | 81 | PQ507924 |

**Table S3.** Jumbo phages genome characterization. Table S4 Bacteria Genomes.

**Table S5.** qPCR Primers.

## Supplemental Methods

### Phage and bacteria sequencing

Phage DNA was extracted using the QIAamp Ultrasens Virus Kit (Qiagen). Bacterial DNA was extracted using the DNEasy Blood and Tissue Kit (Qiagen). All DNA libraries were prepared using the Nextera XT Library Preparation Kit (Illumina). Phage genome sequencing was completed using a paired-end approach (2 × 150 bp) on the iSeq100 (Illumina), and genomes were assembled using the de novo assembly algorithm of CLC Genomics Workbench (CLC Genomics, Qiagen, version 9.5.3). Genome sequencing of bacterial isolates was completed using the MiSeq (Illumina), and genomes were assembled using the Genome Assembly Service from the Bacterial and Viral Bioinformatics Research Center (BV-BRC version 3.36.16.3).

### Phage Genome Annotations

Phage genomes were annotated using the Genome Annotation Service from the Bacterial and Viral Bioinformatics Research Center (BV-BRC version 3.36.16.3) to determine the number of coding sequences (CDS).

Conserved domains were found using the Conserved Domain Database from NCBI. Phage proteomic tree and genome alignments were produced using ViPTree (version 4.0).

### TEM Imaging

A carbon-coated grid (PELCO SynapTek Grids, product# 01754-F) was placed on a drop of 10 µL of freshly made phage stock. The grids were negatively stained with 2% uranyl acetate (pH 4.0) for 45 seconds. Imaging was performed using Jeol 1400 plus located at the University of California, San Diego—Cellular and Molecular Medicine Electron Microscopy Core (RRID: SCR_022039).

### Live-cell fluorescence microscopy

*E. cloacae* cells (4 μL of overnight culture) were inoculated on imaging pads (1% agarose, 25% LB) in welled microscope slides. Slides were incubated at 37 °C for 2 h in a humid chamber to allow for cell growth. 8 μL of phage lysate was added to the pads and infection was allowed to proceed at 37 °C in the humid chamber. At the appropriate time point, imaging pads were stained with 8 μl of dye mix (3 μg/ml FM4-64, 2 μg/ml DAPI) and imaged. Samples were imaged using the DeltaVision Elite deconvolution microscope (Applied Precision). Images were deconvolved using the conservative algorithm in the DeltaVision softWoRx program-v5.5.1 and processed in ImageJ.

## Abbreviations

Si: Silp
Ca: Carethers
Re: Rett
Ar: Arina
Sa: Sayo
CL: Chasing Life
WE: Winchester Ellie
BP: Big Picture

